# Selectivity profile of the poly(ADP-ribose) polymerase (PARP) inhibitor, A-966492

**DOI:** 10.1101/119818

**Authors:** Ann-Gerd Thorsell, Herwig Schüler

## Abstract

Among the clinical inhibitors of poly(ADP-ribose) polymerases (PARPs) and the commonly used PARP research tool compounds, veliparib and niraparib were recently identified as the most selective inhibitors of PARP1 and PARP2. We characterized the potency of A-966492, a PARP inhibitor with a chemical structure similar to veliparib and niraparib, in *in vitro* inhibition experiments of six PARP family enzymes. We find that the selectivity of A-966492 for PARP1 and PARP2 is intermediate between veliparib and niraparib.

## Introduction

Poly(ADP-ribose) polymerases (PARPs) use NAD^+^ as co-substrate to ADP-ribosylate their target proteins. PARP activities contribute to diverse cellular events including DNA repair, control of transcription, and protein localization, maturation, and degradation. PARP1 continues to be an important target for cancer drug development and two compounds, Lynparza (olaparib; AZD2281; KU0059436) and Rubraca (rucaparib; AG-014699), have been approved by the U.S. Federal Drug Administration. Neither olaparib nor rucaparib are selective inhibitors of PARP1; both affect the activities of the closely related PARP2 and several other PARP family enzymes.^5, 6^ In order to improve PARP inhibitor based therapies, it is important to understand the individual functions of the diverse PARP family members, and the effect of inhibiting them in a disease relevant context.^6, 7^ Thus, research outcomes need to be interpreted in relation to the selectivity of the particular enzyme inhibitor used.

We recently published a selectivity profiling study of PARP and tankyrase inhibitors,^4^ and determined that veliparib (ABT-888) and niraparib (MK-4827) had the highest selectivity for PARP1 and PARP2 of the compounds tested.^4^ Here, we report a profiling analysis of a similar compound, A-966492, carried out under identical conditions.

## Materials and Methods

A-966492 was purchased from Selleck. All other reagents, all proteins and methods were as previously described.^4^ Enzymatic inhibition assays were carries out using technical replicates, and repeated 1-4 times.

## Results and Discussion

The PARP inhibitor A-966492 has a chemical structure similar to veliparib and niraparib (**Figure 1A**).^2^ A-966492 is well characterized, and a crystal structure of a very similar derivative of the same scaffold (**Figure 1B**; PDB entry 3L3M) implies that its binding mode is known.^2^ Consequently, it was important to characterize the potency of A-966492 under the conditions used in our recent PARP inhibitor profiling analysis.^4^

**Figure 1:**
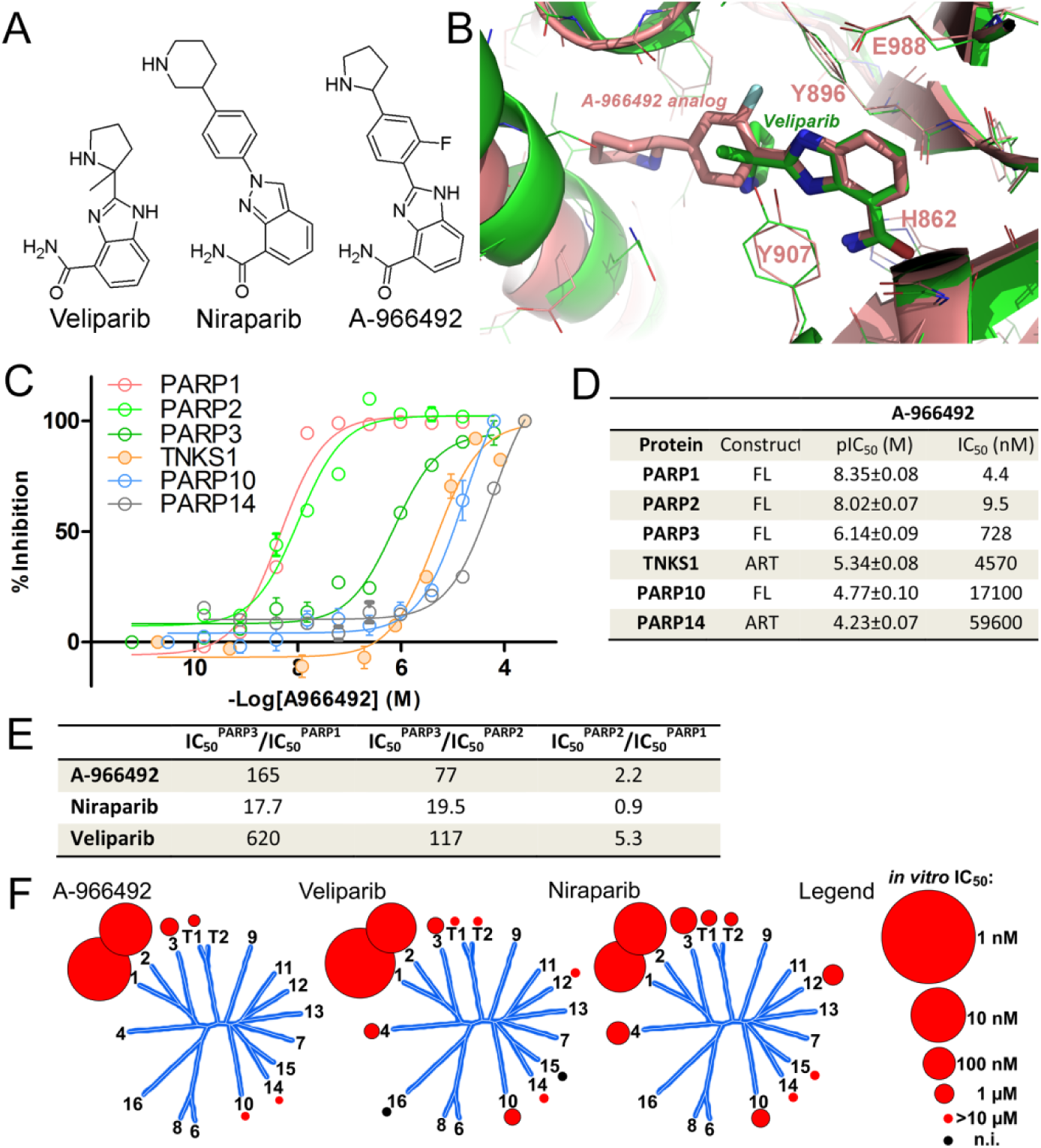
A-966492 is a potent and selective inhibitor of PARP1 and PARP2. (**A**) Chemical structures of veliparib, niraparib, and A-966492. (**B**) Binding modes of veliparib (in green; PDB: 3KJD)^1^ and the piperidine analogue of A-966492 (in salmon; PDB: 3L3M).^2^ PARP1 side chains of the HYYE motif^3^ are indicated. (**C**) Concentration response curves for A-966492. (**D**) Potencies calculated from the data shown in panel C. (**E**) Selectivity ratios for the three most PARP1- and PARP2-selective PARP inhibitors. (**F**) Schematic representation of PARP inhibitor potencies mapped on a phylogenetic tree of human PARP enzymes. Red sphere sizes are proportional, on a logarithmic scale, to IC_50_ values in the range 1 nM - 10 μM; red dots indicate IC_50_ values higher than 10 μM; black dots indicate no inhibition detected.

We determined IC_50_ values of A-966492 mediated inhibition of six PARP family enzymes in *in vitro* enzymatic activity assays (**Figure 1C,D**). The results show that A-966492 is a potent inhibitor of PARP1 and PARP2. IC50 values are in the nanomolar range; likely due to different methods used we determined somewhat higher values than those reported previously.^2^ Our results also show that A-966492 has considerable selectivity over PARP3, TNKS1, and the mono-ADP-ribosyltrasferases, PARP10 and PARP14 (**Figure 1C,D**). The selectivity ratios for PARP1 and PARP2 over PARP3, the next nearest relative, are higher than the corresponding ratios for niraparib, but not as high as for veliparib (**Figure 1E**). We conclude that, of the compounds tested under our conditions, veliparib is the most PARP1 and -2 selective, followed by A-966492, followed by niraparib (**Figure 1F**).

## Author Contributions

A.-G.T. designed research, carried out experiments, and analyzed data; H.S. analyzed data and wrote the manuscript.

## Notes

The authors declare no competing financial interest.

## Acknowledgments

This work was financed by the IngaBritt och Arne Lundbergs Research Foundation, the Swedish Cancer Society, and the Swedish Research Council

